# Increased regional P2X7R uptake detected by [^18^F]GSK1482160 PET in a tauopathy mouse model

**DOI:** 10.1101/2024.01.27.575823

**Authors:** Yanyan Kong, Lei Cao, Jiao Wang, Junyi Zhuang, Yongshan Liu, Lei Bi, Yifan Qiu, Yuyi Hou, Qi Huang, Fang Xie, Yunhao Yang, Kuangyu Shi, Axel Rominger, Yihui Guan, Hongjun Jin, Ruiqing Ni

**Affiliations:** PET Center, Huashan Hospital, Fudan University, Shanghai, China; Institute for Regenerative Medicine, University of Zurich, Zurich, Switzerland; Lab of Molecular Neural Biology, School of Life Sciences, Shanghai University, Shanghai, China; Guangdong Provincial Engineering Research Center of Molecular Imaging, the Fifth Affiliated Hospital, Sun Yat-Sen University, Zhuhai, 519000, Guangdong Province, China; Department of Nuclear Medicine, University Hospital, Inselspital Bern, Bern, Switzerland; Institute for Biomedical Engineering, University of Zurich & ETH Zurich, Zurich, Switzerland

**Author notes:** Corresponding author: Hongjun Jin, Address: 52 E, Meihua Road, Zhuhai 519000, China, Ruiqing Ni,; Address: Wagistrasse 12, 9^th^ floor, Zurich 8952 Switzerland.

**Keywords:** Alzheimer’s disease, amyloid-beta, glia, P2X7R, PET, tau

## Abstract

Neuroinflammation plays an important role in Alzheimer’s disease and primary tauopathies. The aim of the current study was to map [^18^F]GSK1482160 for imaging of purinergic P2X7R in Alzheimer’s disease and primary tauopathy mouse models. MicroPET was performed using [^18^F]GSK1482160 in widely used mouse models of Alzheimer’s disease (APP/PS1, 5×FAD and 3×Tg), 4-repeat tauopathy (rTg4510) mice and age-matched wild-type mice. Increased uptake of [^18^F]GSK1482160 was observed in the cortex and basal forebrain of 7-month-old rTg4510 mice compared to age-matched wild-type mice and compared to 3-month-old rTg4510 mice. Nonparametric Spearman’s rank analysis revealed a positive correlation between tau [^18^F]APN-1607 uptake and [^18^F]GSK1482160 in the hippocampus of rTg4510 mice. No significant differences in the uptake of [^18^F]GSK1482160 were observed between wild-type mice and APP/PS1 mice (5, 10 months), 5×FAD mice (3, 7 months) or 3×Tg mice (10 months). Immunofluorescence staining further indicated the distribution of P2X7Rs in the brains of 7-month-old rTg4510 mice with accumulation of tau inclusion compared to wild-type mice. These findings provide in vivo imaging evidence for increased P2X7R in the brains of tauopathy model mice.

## Introduction

Alzheimer’s disease (AD) is the most common cause of dementia and is pathologically characterized by amyloid-beta (Aβ) plaque and tau tangle deposition (56). Primary tauopathies such as frontotemporal lobar degeneration (FTLD), corticobasal degeneration (CBD) and progressive supranuclear palsy (PSP) are pathologically characterized by tau inclusions. Neuroinflammation plays an important role in AD and primary tauopathies (17, 19, 48). Purinergic signalling occurs in individuals with cognitive disturbances (22), cognitive impairment and neuropsychiatric symptoms of AD (51). Purinergic receptors, particularly P2X7Rs, play important roles in chronic immune and inflammatory responses (2, 11). In the central nervous system, P2X7R is expressed on microglia, astrocytes (2) and oligodendrocytes, but its expression on neurons is still unclear (11). P2X7R has proinflammatory functions by increasing the level of adenosine triphosphate, which activates P2X7R and leads to proinflammatory cytokines. Activation of P2X7Rs contributes to NOD-, LRR- and pyrin domain-containing protein 3 (NLRP3) inflammasome activation and the secretion of mature interleukin (IL)-1β (35, 49, 54). Increased expression and activation of P2X7Rs have been reported in postmortem brains from patients with AD, FTLD, or PSP (5, 40). Polymorphisms of the *p2rx7* gene impact the risk of developing AD (55). In addition, P2X7Rs are involved in important pathological processes involved in the development of AD, including Aβ production and plaque formation, tau tangles, oxidative stress, and chronic neuroinflammation. There is a vicious cycle between Aβ and alterations in the levels of P2X7Rs: P2X7Rs promote proinflammatory pathways via Aβ-mediated chemokine release (37), particularly through the chemokine CCL3, which is associated with pathogenic CD8+ T-cell recruitment (37). In addition, P2X7R affects Aβ production by influencing alpha-secretase-dependent amyloid precursor protein (APP) processing (9, 41) and increasing plaque formation mediated by glycogen synthase kinase-3β (12). P2X7R also influences the aggregate burden of tau in human tauopathies and results in distinct signalling in microglia and astrocytes (1, 2).

Upregulated expression levels of P2X7Rs in AD mouse models have been reported, such as in rats injected with Aβ_42_ (40), APP/PS1 mice (35, 37), J20 mice (38) and tauopathy THY-Tau22 mice (5); moreover, deficiency of the *p2rx7* gene or pharmacological inhibition of P2RX7 has been shown to improve plasticity and cognitive ability in P301S tau transgenic mice (10, 52) and amyloidosis mice (37). Pharmacological or genetic P2X7R blockage reversed the proteasomal impairment induced by P301S tau in mice (4). Thus, P2X7R is a promising target for neuroinflammation imaging and a therapeutic target for AD and primary tauopathies.

Several PET tracers for P2X7R have been developed (64), including [^11^C]/[^18^F]GSK1482160 (20, 57), [^18^F]4A (21), [^11^C]SWM139 (26), [^11^C]A-740003, [^18^F]JNJ-64413739 (8), [^123^I]TZ6019 (28), [^18^F]FTTM (14), and [^18^F]IUR-1601 (15). Increased levels of [^11^C]GSK1482160 (57) have been detected in the cortex and hippocampus of mice treated with lipopolysaccharide in a blockable manner (57). To date, the in vivo pattern of P2X7R in AD models and in tauopathy models has not been elucidated, with only one study reporting on a 12- to 13-month-old APP/PS1 mouse model generated from [^18^F]4A (21). An earlier autoradiography study using [^11^C]SMW139 showed comparable binding in temporal cortex tissue from AD patients and controls (26).

The aim of the current study was to evaluate changes in the levels of P2X7R by using [^18^F]GSK1482160 PET in widely used mouse models of AD and 4-repeat tauopathy. We assessed the alterations of P2X7R in APP/PS1 mice (5, 10 months), 3×Tg mice (10 months), 5×FAD mice (3, 7 months), rTg4510 mice (3, 7 months) and age-matched wild-type mice. Ex vivo staining was performed to determine the distribution of P2X7R in the brain of the mice underwent in vivo imaging.

## Methods

### Animal models

The animal models used in the study are summarized in **Table 1**. Sixty mice were included in total, including rTg4510 mice [STOCK Tg(Camk2a-tTA)1Mmay Fgf14Tg(tetO-MAPT*P301L)4510Kha/J] (Jax Laboratory) (53) and APP/PS1 mice [B6. Cg-Tg(APPswe,PSEN1dE9)85Dbo/Mmjax] (25), 3×Tg mice [B6;129-Psen1tm1MpmTg(APPSwe, tauP301L)1Lfa/Mmjax] (Jax Laboratory) (45), and 5×FAD mice [B6. Cg-Tg(APPSwFlLon,PSEN1*M146L*L286V)6799Vas/Mmjax] (Jax Laboratory) (44). Wild-type C57BL6 mice were obtained from Charles River, Germany, and Cavins Laboratory Animal Co., Ltd., of Changzhou. Mice were housed in ventilated cages inside a temperature-controlled room under a 12-h dark/light cycle. Pelleted food and water were provided ad libitum. Paper tissue and red mouse house shelters were placed in cages for environmental enrichment. The in vivo PET imaging and experimental protocol were approved by the Institutional Animal Care and Ethics Committee of Huashan Hospital of Fudan University and Sun Yat-sen University and performed in accordance with the National Research Council’s Guide for the Care and Use of Laboratory Animals. All experiments were carried out in compliance with national laws for animal experimentation and were approved by the Animal Ethics Committee of Fudan University and Sun Yat-Sen University.

**Table 1.** Information on the animal models used in the study.

| Animal model | Age (month) | [<sup>18</sup>F]GSK1482160 PET | [<sup>18</sup>F]APN-1607 PET |
| --- | --- | --- | --- |
| WT | 3 | 4 M |  |
|  | 5 | 5 M |  |
|  | 7 | 5 M |  |
|  | 10 | 5 M |  |
| rTg4510 | 3 | 6 M | 5 M |
|  | 7 | 8 M | 6 M |
| APP/PS1 | 5 | 6 M |  |
|  | 10 | 6 M |  |
| 3×Tg | 10 | 3 M |  |
| 5×FAD | 3 | 3 M/3 F |  |
|  | 7 | 3 M/3 F |  |

M: male, F: female. Sixty mice were included in total. Detailed [<sup>18</sup>F]APN-1607 PET SUVR data can be found in (32).

### Radiosynthesis

[^18^F]GSK1482160 was synthesized and radiolabelled based on nucleophilic aliphatic substitution according to protocols described previously (20, 62). The identities of the final products were confirmed by comparison with the high-performance liquid chromatography (HPLC) retention times of the nonradioactive reference compounds obtained by coinjection using a Luna 5 μm C18(2) 100 Å (250 mm×4.6 mm) column (Phenomenex) with acetonitrile and water (60:40) as the solvent and a 1.0 mL/min flow rate. A radiochemical purity > 95% was achieved for [^18^F]GSK1482160 (molar activity, 1.48 GBq/ml) (21, 62) (the HPLC results are shown in **SFig. 1**).

### MicroPET

PET experiments were performed using a Siemens Inveon PET/computed tomography (CT) system (Siemens Medical Solutions, Knoxville, United States) (30). Prior to the scans, the mice were anaesthetized using isoflurane (2.0–3.0%) in medical oxygen (1 L/min) at room temperature with an isoflurane vaporizer (Molecular Imaging Products Company, United States). The mice were positioned in a spread-up position on the imaging bed and subjected to inhalation of the anaesthetic (1.5–2.5% isoflurane) during the PET/CT procedure. A single dose of [^18^F]GSK1482160 (∼0.37 MBq/g body weight, 0.1–0.2 mL) was injected into the animals through the tail vein under isoflurane anaesthesia. Static PET/CT images were obtained 10 min after intravenous administration of [^18^F]GSK1482160 at 50-60 min. PET/CT images were reconstructed using the ordered subset expectation maximization 3D algorithm (OSEM3D), with a matrix size of 128×128×159 and a voxel size of 0.815 mm×0.815 mm×0.796 mm. The data were reviewed using Inveon Research Workplace (IRW) software (Siemens, United States). Attenuation corrections derived from hybrid CT data were applied.

### Imaging Data Analysis

The images were processed and analysed using PMOD 4.4 software (PMOD Technologies Ltd., Zurich, Switzerland). The time-activity curves were deduced from specific volumes of interest that were defined based on mouse magnetic resonance imaging T_2_-weighted images and the Ma-Benveniste-Mirrione atlas (in PMOD). Radioactivity is presented as the standardized uptake value (SUV) (decay-corrected radioactivity per cm^3^ divided by the injected dose per gram body weight). The brain regional SUVRs were calculated using the cerebellum (Cb) as the reference region as described in an earlier PET study using [^18^F]JNJ-64413739 in rodents (3). A mask was applied for signals outside the brain volumes of interest for illustration. To understand the link between tau deposits and P2X7R alterations in the brain, we performed non-parametric Spearman’s correlation analysis between the uptake of [^18^F]GSK1482160 and the tau tracer [^18^F]APN-1607 in the brain of rTg4510 mice. Data for [^18^F]APN-1607 SUVR (with Cb as the reference region) in the same rTg4510 mice underwent [^18^F]GSK1482160 imaging was obtained from an earlier study (n = 11) (32).

### Immunohistochemistry

After in vivo imaging, the mice were anaesthetized with tribromoethanol, perfused with ice-cold 0.1 M phosphate-buffered saline (PBS, pH 7.4) and 4% paraformaldehyde in 0.1 M PBS (pH 7.4), fixed for 36 h in 4% paraformaldehyde (pH 7.4) and subsequently stored in 0.1 M PBS (pH 7.4) at 4°C. The brain was placed in 30% sucrose in PBS until it sank. The brain was embedded in OCT gel (Tissue-Tek O.C.T., Sakura, USA). Coronal brain sections (20 μm) were cut around the bregma 0 to -2 mm using a Leica CM1950 cryostat (Leica Biosystems, Germany). For P2X7R immunofluorescence labelling, sections were blocked in blocking buffer containing 3% bovine serum albumin (BSA), 0.4% Triton X-100, and 5% normal goat serum (NGS) in PBS for 2 h at room temperature. After washing with PBS for 3×10 minutes, the sections were incubated with primary antibodies in blocking buffer overnight at 4°C, incubated with donkey anti-rat IgG H&L (Alexa Fluor® 647) (1:500, ab150155, Abcam) in blocking buffer for 2 hours at room temperature and subsequently washed with PBS for 3×10 minutes.

For Aβ and tau staining, coronal brain sections (3 μm) were cut using a Leica RM2016 Microtome (Leica, Germany). The sections were first washed in PBS 3×10 minutes, followed by antigen retrieval for 20 minutes in citrate buffer (pH 6.0) at room temperature. After antigen retrieval in citrate buffer at room temperature, the sections were permeabilized and blocked in 3% bovine serum albumin for 30 minutes at room temperature with mild shaking. Paraffine-embedded sections were incubated overnight at 4°C with primary antibodies against Aβ and phospho-Tau (Ser202, Thr205), as described earlier (29). The next day, the slices were washed with PBS 3×5 minutes, incubated with secondary antibody for 2 hours at room temperature and washed 3×5 minutes with PBS. The sections were incubated for 10 minutes in 4’,6-diamidino-2-phenylindole (DAPI) at room temperature and mounted with antifade mounting media (31). The brain sections were imaged at ×20 magnification using a Pannoramic MIDI slide scanner (3DHISTECH) using the same acquisition settings for all brain slices. The images were analysed by using ImageJ (NIH, U.S.A.).

### Statistics

Two-way ANOVA with Sidak’s post hoc analysis was used for comparisons between groups (GraphPad Prism 9.0, CA, USA). Nonparametric Spearman’s rank correlation analysis was used to analyse the associations between the regional SUVRs of [^18^F]GSK1482160 and [^18^F]APN-1607. A p value less than 0.05 was considered to indicate statistical significance. The data are shown as the mean ± standard deviation.

## Results

### The regional uptake of [^18^F]GSK1482160 was greater in rTg4510 mice than in age-matched wild-type mice and correlated with tau deposits in the hippocampus

The HPLC and quality control results for [^18^F]GSK1482160 are shown in **SFig. 1**. The dynamics of [^18^F]GSK1482160 in the periphery of mouse and rat models and the in vitro and in vivo stabilities of [^18^F]GSK1482160 were demonstrated in a recent study (62). We found that [^18^F]GSK1482160 exhibited relatively lower uptake inside the mouse brain than outside the mouse brain (**SFig. 2**). In contrast to translocator protein (TSPO) imaging, which has a high cerebellar update rate (7), we found that the cerebellar uptake of [^18^F]GSK1482160 was not greater than that of cerebral regions. In an earlier P2X7R imaging study using [^18^F]JNJ-64413739 in a rodent model, the cerebellum was also used as the reference region for SUVR calculations (3). The cerebellum is suitable for use as a reference brain region for [^18^F]GSK1482160.

rTg4510 mice are known to develop tau pathology at approximately 5 months of age (53). To determine whether and when alterations in the levels of P2X7R in the brain of the rTg4510 tau mouse model are related to tau accumulation, we assessed [^18^F]GSK1482160 imaging in 3- and 7-month-old (with tau pathology) rTg4510 mice. We quantified the regional [^18^F]GSK1482160 SUVR using the cerebellum as a reference region. We observed that the [^18^F]GSK1482160 SUVR was greater in the cortex and basal forebrain system of 7-month-old rTg4510 mice than in age-matched wild-type mice (**Figs. 1b, f, g**) and was greater in the cortex, basal forebrain system, striatum and amygdala of 7-month-old rTg4510 mice than in 3-month-old rTg4510 mice (**Figs. 1e-g**).

**Fig. 1.**
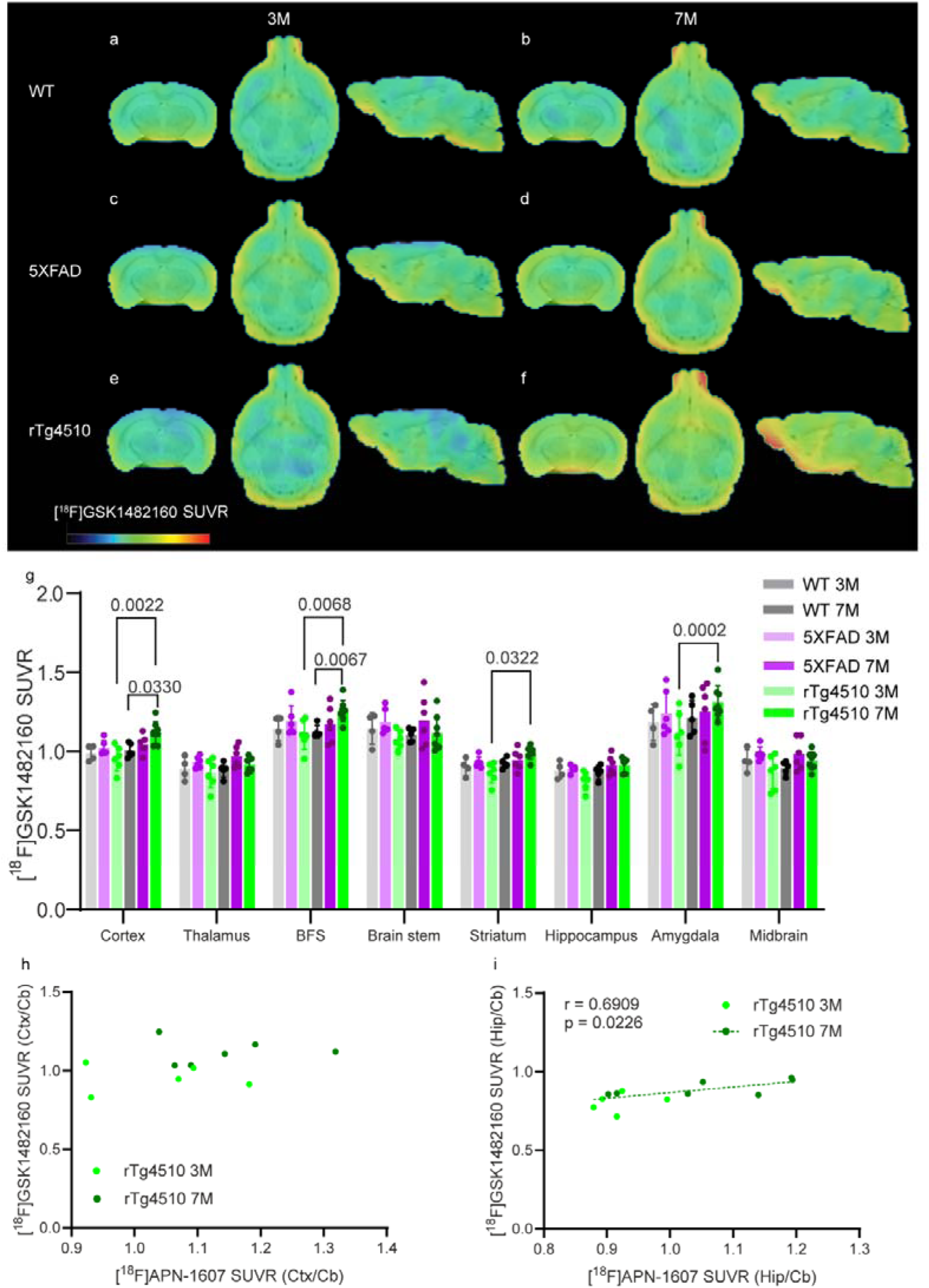
Increased regional [^18^F]GSK-1482160 brain uptake in 7-month-old rTg4510 mice compared to age-matched wild-type mice and correlation with [^18^F]APN-1607 uptake in the brain. **(a-f)** Representative [^18^F]GSK-1482160 SUVR images of 3- and 7-month-old WT (a, b), 5×FAD (c, d), and rTg4510 (e, f) mice; SUVR scale, 0-2.2. BFS, basal forebrain system; (g) Quantification of [^18^F]GSK-1482160 SUVRs in WT, 5×FAD and rTg4510 mice. **(h, i)** Non-parametric Spearman’s rank analysis of [^18^F]GSK-1482160 and [^18^F]APN-1607 SUVR (Cb as the reference region) in rTg4510 mice (3-month n = 5, 7-month, n = 6).

To understand the link between tau accumulation and P2X7R alterations in the brain, we performed correlation analysis between [^18^F]GSK1482160 and the tau tracer [^18^F]APN-1607 in eleven rTg4510 mice (including five 3 months-of-age and six 7 months-of-age). The description for radiosynthesis and mciroPET method for [^18^F]APN-1607 imaging in rTg4510 mice and wild-type mice can be found in (32). The data for [^18^F]APN-1607 SUVR (with Cb as the reference region) in the same rTg4510 mice were obtained from our recent study (32). Nonparametric Spearman’s rank analysis revealed a positive correlation between tau [^18^F]APN-1607 uptake and [^18^F]GSK1482160 in the hippocampus of rTg4510 mice (r = 0.6909, p = 0.0226, n = 11, **Fig. 1i**)

### The uptake of [^18^F]GSK1482160 did not differ between APP/PS1, 5×FAD and 3×Tg mice and age-matched wild-type mice

Next, we assessed the changes in the level of P2X7R in the brains of mouse models of AD amyloidosis. 5×FAD mice develop Aβ plaques at approximately 5 months of age. Here, we chose 3 months and 7 months as the pre- and post-pathological time points, respectively. Here, we found that there was no difference in the [^18^F]GSK1482160 SUVR in the brains of 5×FAD mice (3-. 7-months) compared to age-matched wild-type mice (**Figs. 1a-d, g**).

APP/PS1 mice develop Aβ plaques at approximately 6 months of age. 3×Tg mice are models of both cerebral Aβ plaques and tau pathology; however, due to genetic drift, recent studies from our group and other groups have shown limited pathology at the age of 10 months (27, 45). Here, we chose APP/PS1 mice at 5 and 10 months of age and 3×Tg mice at 10 months of age for [^18^F]GSK1482160 imaging. We observed no difference in [^18^F]GSK1482160 SUVR in other mouse models of APP/PS1 (5, 10 months) (**Fig. 2**) or in 3×Tg mice (10 months) (**Fig. 2**).

**Fig. 2.**
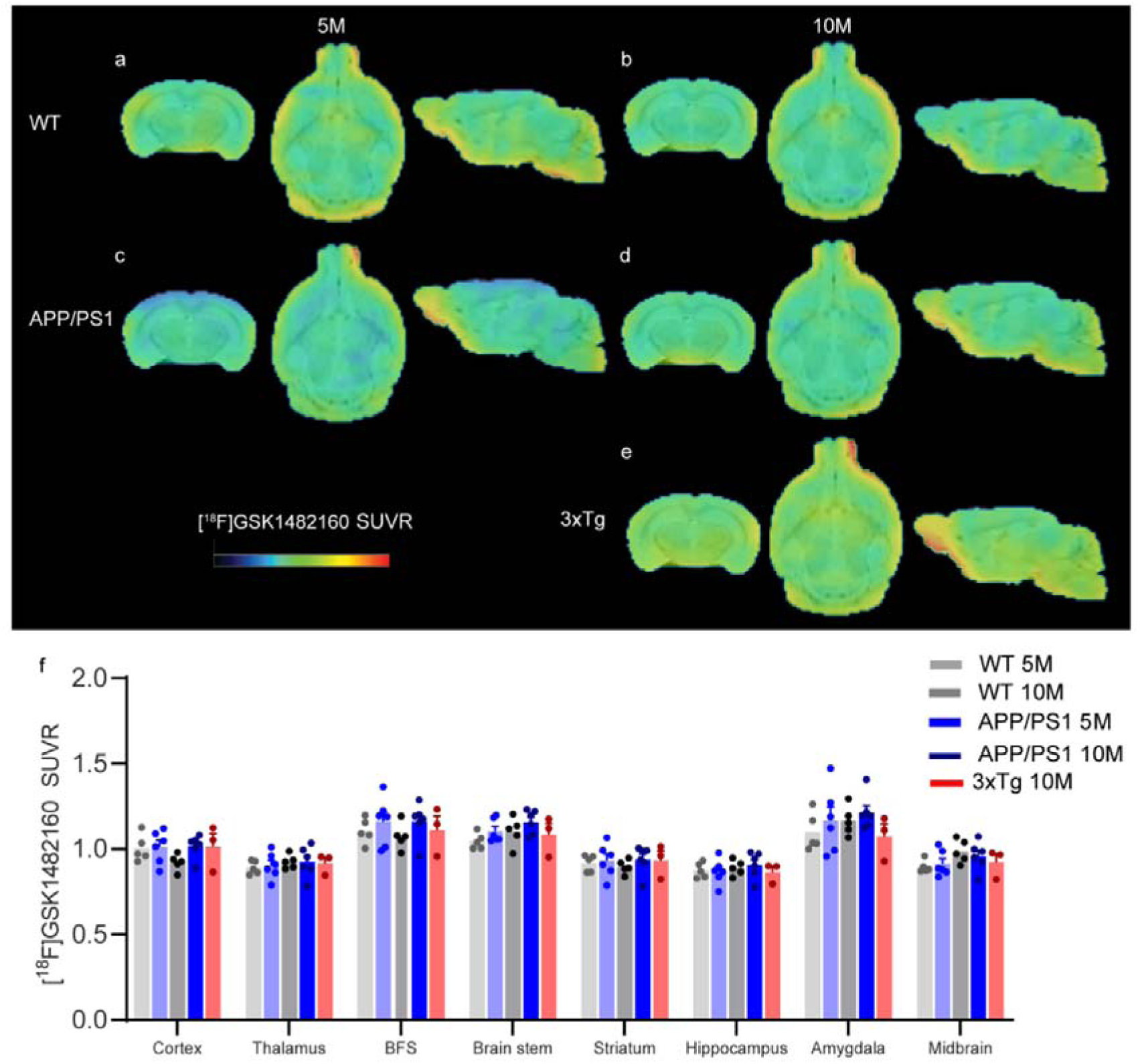
No difference in regional [^18^F]GSK-1482160 expression between 5- or 10-month-old APP/PS1 mice or 3×Tg mice and wt mice. (**a-e**) Representative [^18^F]GSK-1482160 SUVR images of 5- and 10-month-old WT (a, b), APP/PS1 (c, d), and 10-month-old 3×Tg (e) mice; SUVR scale 0- 2.2 BFS, basal forebrain system; (**f**) Quantification of [^18^F]GSK-1482160 SUVRs in WT, APP/PS1 mice and 3×Tg mice.

### P2X7R distribution in the mouse brain

Ex vivo immunofluorescence staining was performed on the mouse brain tissue slices after in vivo imaging. The P2X7R signal in the brains of APP/PS1 mice and WT mice appeared comparable. The accumulation of tau inclusion was validated by using phospho-Tau staining of brain tissue slices from 7-month-old rTg4510 mice and WT mice (**Figs. 3a-c**). Tau tangles and were detected in the cortex and hippocampus, of 7-month-old rTg4510 mice. Amyloid-beta deposits were detected in the cortex, hippocampus and thalamus of both 5×FAD mice at 7 months of age (**Figs. 3d-f**) and 10-month-old APP/PS1 mice at 10 months of age (**Figs. 3g-i**). The P2X7R signal in the brains of the rTg4510 mice was higher in the hippocampus than in the cortex based on the ex vivo staining, which was slightly different from the in vivo pattern (**Figs. 4a-l**).

**Fig 3.**
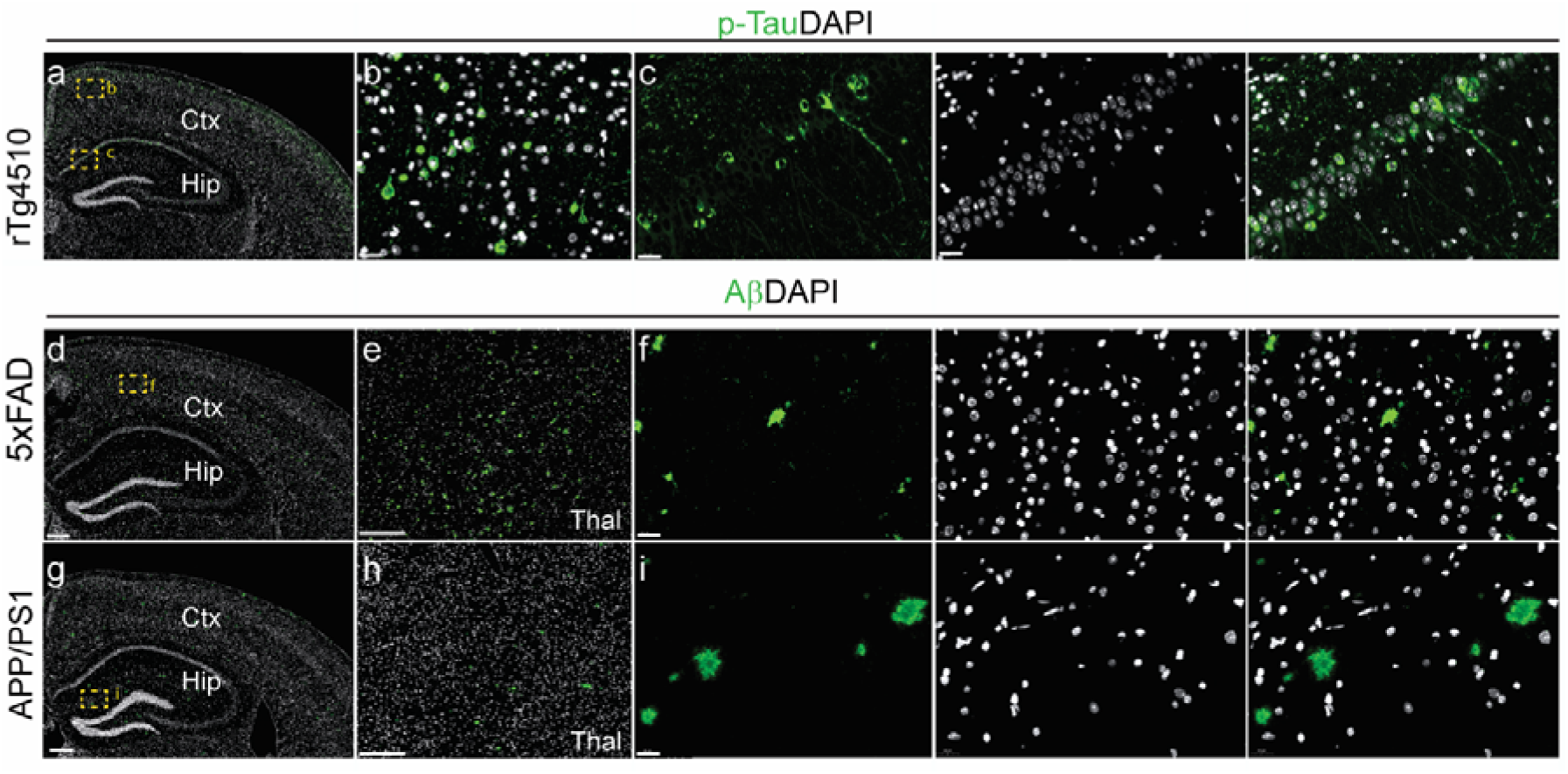
Staining of tau inclusions in the brain from 7 month-old rTg4510 mice and amyloid-beta deposits in the brains of 7-month-old 5×FAD mice and 10-month-old APP/PS1 mice. (**a-c**) An overview and zoomed-in view of the staining of phospho-Tau (p-Tau, green) in coronal brain slices from 7-month-old rTg4510 mice. (**c**) Zoomed-in view showed tau tangles in the cortex (Ctx), and hippocampus (Hip). (**d-i**) Overviews and zoomed-in view of the staining of coronal brain slices from 7-month-old 5×FAD mice and 10-month-old APP/PS1 mice. Zoomed-in views revealing the presence of amyloid-beta deposits (green) in the cortex (Ctx) of 5×FAD mice and hippocampus (Hip) of 10-month-old APP/PS1 mice (j-l). The yellow squares in the overviews indicate the locations of the zoom-ins. The nuclei were counterstained with DAPI (blue). Scale bar = 200 microns (a, d, e, g, h), or 20 microns (b, c, f, i). Thal, thalamus.

**Fig. 4.**
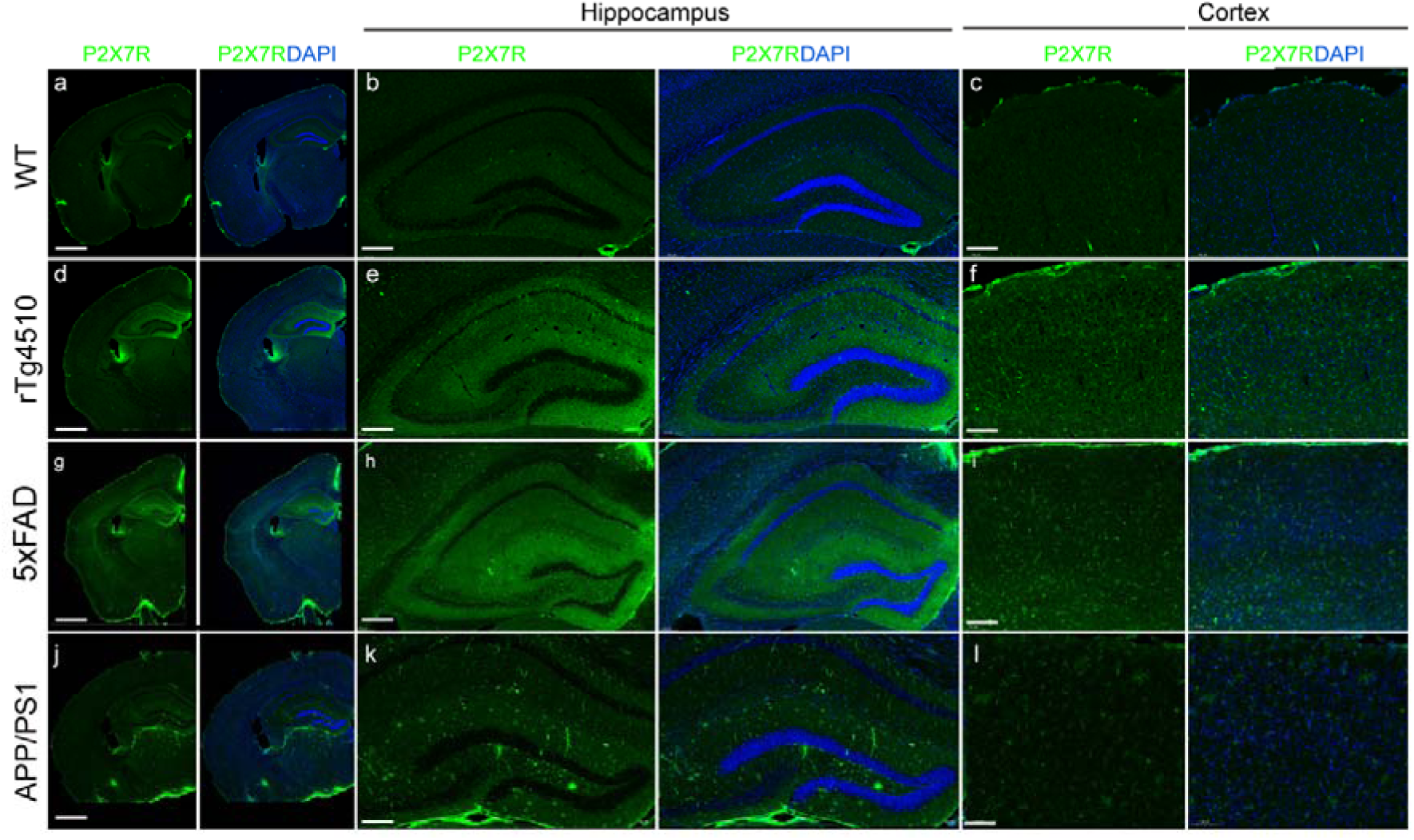
Representative P2X7R staining of coronal brain slices from WT, rTg4510, 5×FAD and APP/PS1 mice. (**a, d, g, j)** Overview and **(b, c, e, f, h, I, k, l**) Zoomed-in view revealing the distribution of P2X7Rs (green) in the hippocampus (Hip) and cortex (Ctx) of 7-month-old WT (a-c), 7-month-old rTg4510 (d-f), 7-month-old 5×FAD (g-i) and 10-month-old APP/PS1 mice. Nuclei were counterstained with DAPI (blue). Scale bar = 1 mm (a, d, g, j), 200 microns (c, f, i, l), 100 microns (b, e, h, k),

## Discussion

Here, we found increased regional [^18^F]GSK1482160 uptake in the cortical and subcortical regions of P2X7Rs in 7-month-old rTg4510 mice compared to age-matched controls and 3-month-old rTg4510 mice. No difference in the regional [^18^F]GSK1482160 SUVR was observed in the brains of APP/PS1 mice (5, 10 months), 5×FAD mice (3, 7 months), or 3×Tg mice (10 months) compared to age-matched wild-type mice.

rTg4510 mice develop tau deposits at 4-5 months of age, which can be detected by PET using [^11^C]PBB3 and [^18^F]PM-PBB3 ([^18^F]APN-1607) (23, 42). In vivo imaging using [^18^F]DPA-714 and [^11^C]AC-5216 has shown increases in TSPO (representing microgliosis) along with tau accumulation (13, 23). Microglial NF-κB has been shown to drive tau spreading and toxicity in the brain of rTg4510 mice (61). In addition, genome-wide RNAseq studies and ex vivo staining have demonstrated increased immunoreactivity for the astrocyte and the microglial marker in the brains of rTg4510 mice (6, 23, 50, 61). A recent study showed that the RNA level was increased in the brain of rTg4510 mice at 6 months and that P2X7R influences the tau aggregate burden in human tauopathies and induces distinct signalling in microglia and astrocytes (1). Our observation of an increase in P2X7R by PET using [^18^F]GSK1482160 in tauopathy mouse brain is in line with the existing autoradiographic study using [^123^I]TZ6019 in the brain tissue from 9-month-old PS19 (P301S tau) mice, with an approximately 35% increase compared to age-matched wild-type mice (28). Tau pathology has been shown to epigenetically remodel neuron-glial cross-talk in AD (63).

In contrast to the increase observed in tau rTg4510 mice, we found no clear difference in [^18^F]GSK1482160 uptake in the brains of 3- or 7-month-old 5×FAD mice. Similar observation was reported in an earlier immunostaining study showing the opposite pattern in P2X7R signalling, with an increase in the brains of tauopathy P301S mice and a reduction in the brains of 5×FAD mice in the same study (16). Another study revealed that P2X7R protein expression in the frontal cortex was approximately 150% greater in 9-month-old 5×FAD mice than in 3-month-old 5×FAD mice and was greater than in wild-type mice (24).

For APP/PS1 mice, regional P2X7Rs in the brains of 5-month-old and 10-month-old APP/PS1 mice were comparable to those in age-matched wild-type mice by using [^18^F]GSK1482160. To date, only one in vivo study of [^18^F]4A PET has been performed in a 12- to 13-month-old APP/PS1 mouse model (21), with the highest binding observed in the anterior commissure, putamen, neocortex and medulla, followed by the cerebellum. Despite the increased area under the curve, the SUVRs (relative to the cerebellum) were comparable between 12- to 13-month-old APP/PS1 and wild-type mice (21). An earlier study showed that immunofluorescence staining of P2X7Rs increased by 20% in the cortex and 30% in the hippocampus of 10-month-old APP/PS1 mice compared to wild-type mice (37), while another western blot analysis showed that P2X7Rs were more abundant in the cortex of 6-month-old and 12-month-old APP/PS1 mice compared to nontransgenic littermates (35). For 3×Tg mice, transcriptomic analysis of the levels of *p2rx7* in 2-, 10- and 20-month 3×Tg mice did not reveal a significant difference (34). In the 19-month-old APPswe mouse model, P2X7R levels were increased in hippocampal tissue compared to those in wild-type tissue, as shown by western blot (no quantification presented) (47).

Given the heterogeneity and dynamic changes in glia, further analysis is needed to determine the differences in the profiles of P2X7R in the brains of mouse models generated with other PET tracers (33), such as [^18^F]DPA-714 for TSPO (33) and [^18^F]SMBT-1 or [^11^C]DED for monoamine-oxidase B (MAO-B) (31, 39, 43, 60). In addition to P2X7R, an increase in P2X4R (59) and a reduction in P2Y12R (18, 36) are also implicated in AD, and comparative imaging study using tracers for P2X7R and P2Y12R will be informative. Moreover, earlier profiling and immunostaining studies showed that TSPO upregulation is selective for proinflammatory polarized astrocytes and microglia (46). In TgF344 rats, astrocytic TSPO upregulation occurs before microglial TSPO upregulation in AD (58). For P2X7R, which are expressed on both astrocytes and microglia, further analysis of the sequence of astrocytic and microglial P2X7R upregulation will be informative.

There are several limitations in the current study. First, sex differences in the expression of P2X7Rs in mouse brains have not been studied. Second, the current study was cross-sectional rather than longitudinal. Further longitudinal assessments in the same set of animal models will be informative. Third, the microPET scans are not dynamic; therefore, BPnd readouts were not available. Moreover, P2X7R expression is highly dependent on the activation state of microglia, and further detailed profiling of microglia in relation to in vivo P2X7R tracer uptake is needed.

## Conclusion

These findings provide in vivo imaging evidence for diverse patterns of P2X7R, with increased P2X7R in the widely used rTg4510 model of 4-repeat tauopathy but not in the amyloidosis models 5×FAD, APP/PS1 or 3×Tg mice. [^18^F]GSK1482160 imaging might be useful for the in vivo evaluation of neuroinflammation in neurodegenerative diseases.

## Declaration

### Ethical approval

The in vivo PET imaging and experimental protocol were approved by the Institutional Animal Care and Ethics Committee of Huashan Hospital, Fudan University and Sun Yat-sen University and performed in accordance with the National Research Council’s Guide for the Care and Use of Laboratory Animals. All the experiments were carried out in compliance with ARRIVE guidelines 2.0, the national laws for animal experimentation.

### Authors’ contributions

The study was designed by KY and RN. KY performed the radiolabelling, HPLC, and microPET. LC and RN performed the microPET analysis. YL, YH, YQ, and LB provided support for the PET data analysis. RN wrote the first draft. HJ provided the GSK1482160 precursor and methodology for radiolabelling. All the authors contributed to the revision of the manuscript. All the authors have read and approved the final manuscript.

## Acknowledgements

The authors thank Dr. Jianfei Xiao, PET Center, Huashan Hospital, Fudan University, for assisting with the radiolabelling and HPLC.

