## Supplementary figure 1, Supplementary figure 2 for "Increased regional P2X7R uptake detected by [^18^F]GSK1482160 PET in a tauopathy mouse model"

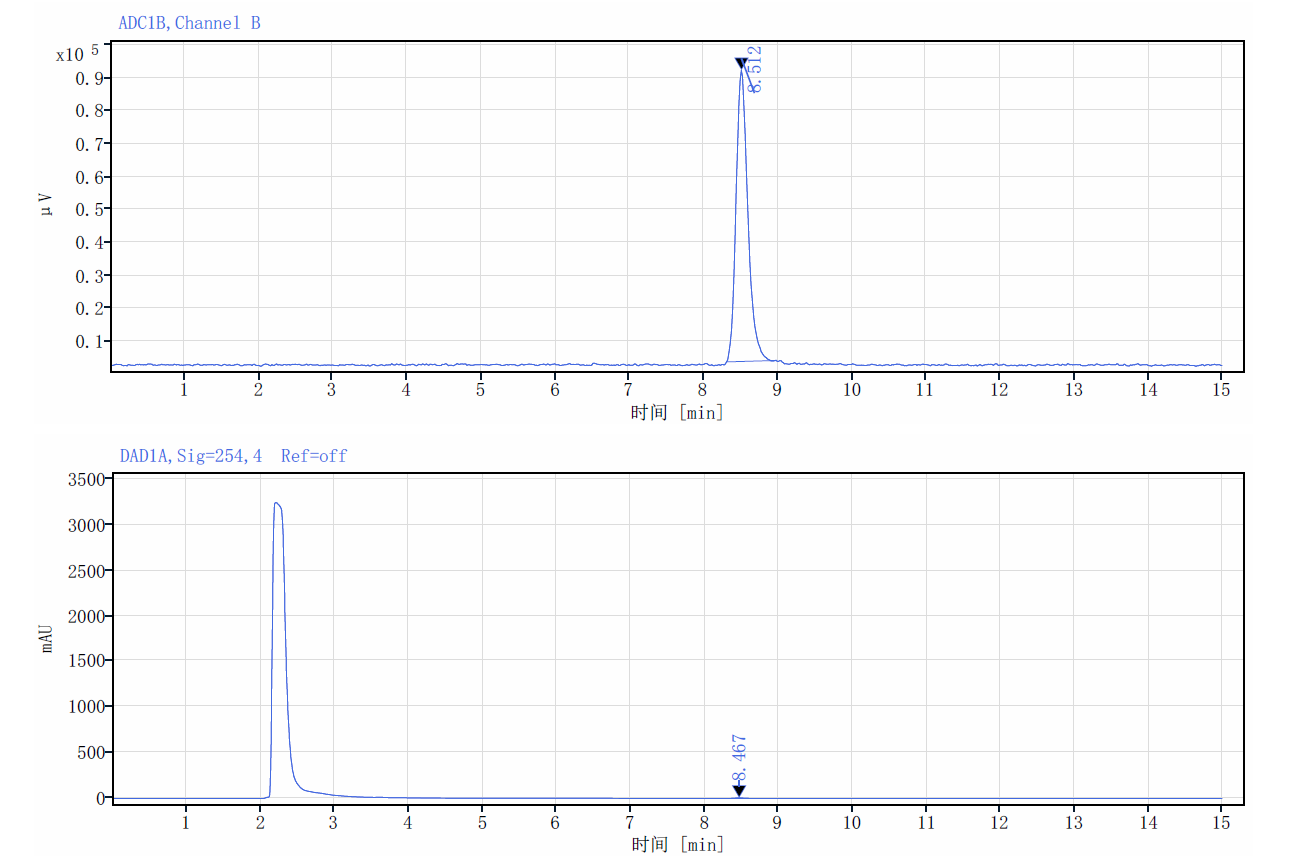

**SFig 1. HPLC chromatogram of synthesized [^18^F]GSK1482160 and the standard.** (**a**) Synthesized GSK1482160 and (**b**) standard.

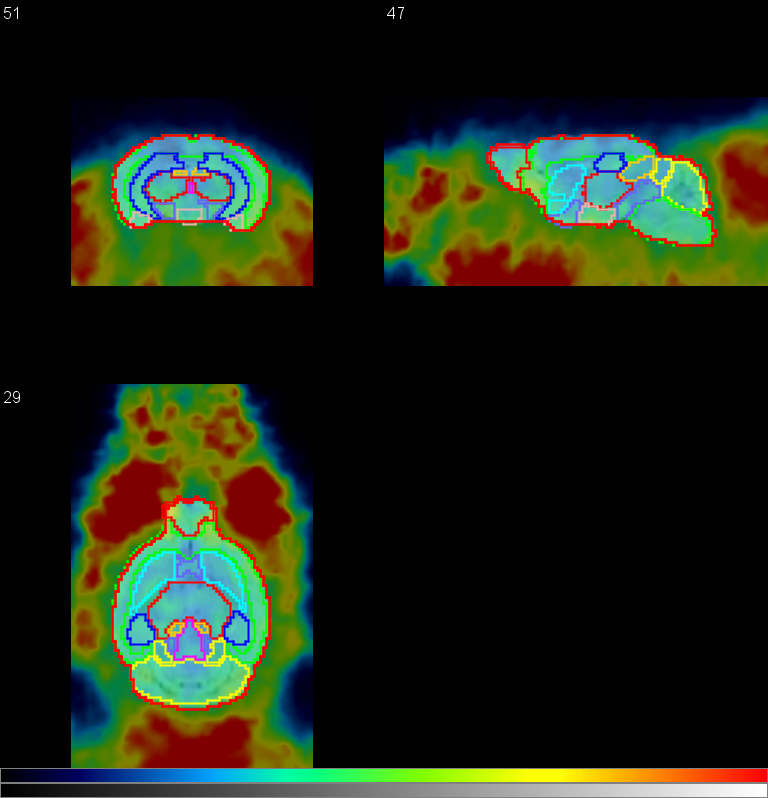

**SFig 2. [^18^F]GSK1482160 exhibited relatively lower brain uptake than did [^18^F]GSK1482160 outside the brain in wild-type mice (averaged 50-60 minutes post injection).** The colour bar indicates an SUVR of 0-2.7. The template used for volume-of interest analysis (overlaid on the images) was the Ma-Benveniste-Mirrione atlas.

**STable 1. Antibodies and reagents used for immunofluorescence staining**

| **Antibodies and reagent** | **Catalog no** | **Dilution** | **Supplier** |
| --- | --- | --- | --- |
| Rat monoclonal anti-P2X7 antibody [1F11] | Ab195356 | 1:200 | Abcam |
| CY3-conjugated goat anti-rat IgG | Gb21302 | 1:300 | Servicebio |
| Anti-Amyloid beta 40, mouse mAb | GB121197 | 1:1200 | Servicebio |
| Goat Anti-Mouse IgG H&L (Alexa Fluor® 488) | ab150113 | 1:500 | Abcam |
| Anti-phospho-TAU (S202/T205), rabbit polyclonal antibody | GB113883 | 1:1000 | Servicebio |
| Alexa Fluor488 goat anti-rabbit IgG | GB25303 | 1:400 | Servicebio |
| CY3-conjugated goat anti-mouse IgG | GB21301 | 1:300 | Servicebio |
| Bovine serum albumin | 36100ES25 | 3% | Yeasen |
| TritonX-100 | X10010 | 0.4% | Abcone |
| Normal goat serum (NGS) | 36119ES03 | 5% | Yeasen |
| 4′,6-diamidino-2-phenylindole (DAPI) | D8200 | 1:2000 | Solarbio |
| Citrate buffer pH 6.0 | G1202 |  | Servicebio |
| Phosphate buffered saline | G0002 |  | Servicebio |
| Anti-fluorescence quenching mounting media | 36307ES08 |  | Yeasen |
